# A skewed picture of functional diversity in global drylands

**DOI:** 10.1101/2024.12.09.627500

**Authors:** Enrico Tordoni, Giacomo Puglielli, Eleonora Beccari, Carlos P. Carmona

## Abstract

Recently, Gross et al. (Nature 632, 808-814) found that plant phenotypic diversity dramatically increases with aridity globally, challenging conventional environmental filtering theories. However, some methodological choices by Gross et al. might overestimate the effect of aridity on phenotypic diversity: (i) We detected a skewed distribution of sampling sites towards the arid end of the analyzed gradient; (ii) We posited that fine-tuning of some statistical parameters might further challenge their estimates. We reanalyzed Gross et al. data accounting for these effects and found a substantially reduced change of phenotypic diversity with increasing aridity. Gross et al. surely represents a cornerstone for future research on trait diversity in a drier world, but their interpretations need to be considered alongside our findings. Importantly, the points raised by our reanalysis provide general guidelines for future research on plant phenotypic diversity changes along environmental gradients.

## Main text

Understanding how aridity shapes plant diversity is crucial in an increasingly drier world. Gross et al.^1^ (hereafter GMB) assessed phenotypic diversity—the volume and integration of plant traits—across a global aridity gradient by sampling sites at different aridity intervals, pooling observations within each interval to estimate the phenotypic diversity at each aridity level. They found that drylands harbour unexpectedly high phenotypic diversity, challenging the idea that harsher environments reduce it^2^. Nevertheless, we believe that their results are influenced by methodological choices, mainly an unbalanced sampling scheme that overrepresents highly arid sites, inflating diversity estimates in the most arid intervals. This bias was amplified by using adaptive bandwidths in trait hypervolume estimations, which further obscure the ecological signal. We reanalysed GMB’s data using a more balanced sampling design and a single common bandwidth and show that the evidence for much higher phenotypic diversity beyond the 0.7 aridity threshold is significantly weaker than suggested by GMB.

The distribution of sites across aridity intervals in GMB is highly unbalanced, with disproportionately more sites in arid regions (see Fig. 1, left panel, in GMB and Extended Data Fig. 1). On average, intervals above the threshold included 2.5 times more sites than those below it (25 vs 10 sites per interval). This uneven sampling resulted in 248 species recorded in the 24 intervals above the threshold compared to just 74 species in the 15 intervals below it (see Supplementary Table 4 in GMB). Furthermore, the larger number of sites above the threshold likely spans a broader set of ecological and biogeographical contexts. Consequently, not only are more species recorded in high aridity intervals, but these species are also likely to exhibit a wider variety of trait combinations.

**Fig 1.**
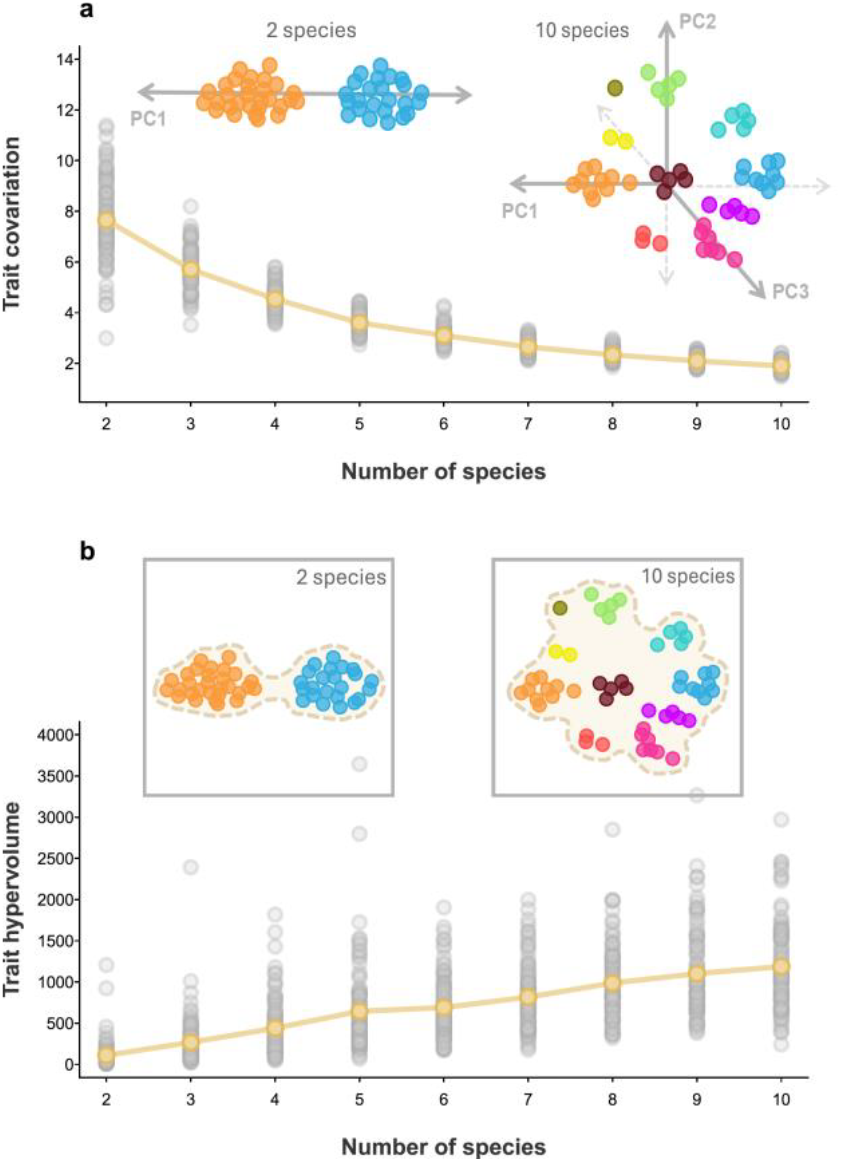
Effect of species richness on trait covariation and hypervolume. **a**, Because individuals of the same species cluster together in trait space, samples including more species are likely to be functionally more complex than species-poor samples. Higher dimensionality is needed to reflect this higher complexity, which is associated with lower trait covariation. The examples show that, for the same number of individuals, a two-species assemblage will generally be well represented by a single dimension of variation, whereas a ten species assemblage requires higher dimensionality. The scatterplot shows the decreasing relationship between trait covariation and species richness in simulated assemblages of 100 individuals (see Supplementary Methods). **b**, Similarly, the number of species in a sample has an important effect on hypervolume size, even when the number of individuals considered is fixed. Due to a sampling effect, samples with two species will tend to occupy a smaller amount of functional space than samples with ten species. The scatterplot shows the increasing relationship between trait hypervolume and species richness in simulated assemblages of 100 individuals.

This unbalanced sampling exacerbated another underlying problem, which is the lack of explicit consideration of species identities. In their analysis, GMB aggregated observations of individual plants from sites within each aridity interval, creating a composite snapshot of plant traits across the gradient. However, individuals of the same species tend to be clustered in the same region of trait space because they exhibit similar trait values^3,4^; overlooking this structure when calculating trait hypervolume and integration introduces a form of pseudoreplication. As a consequence, when samples differ in the number of species, species-rich samples are more likely to include distinct trait combinations than species-poor samples, and hence larger hypervolume and dimensionality estimates (Fig. 1). While differences in species richness are a natural component of variation in functional diversity estimates^5,6^, in the case of GMB these differences largely stem from the unbalanced sampling rather than being a genuine ecological pattern. Because adding an individual of an existing species has a much smaller impact on dimensionality or hypervolume than adding an individual of a new species^7–9^, fixing the number of individuals in the analysis does not address the unbalanced sampling effect. Therefore, trait diversity estimates might be inflated in the most arid intervals, confounding the pattern observed by GMB (Extended Data Figs. 2 and 3).

**Fig 2.**
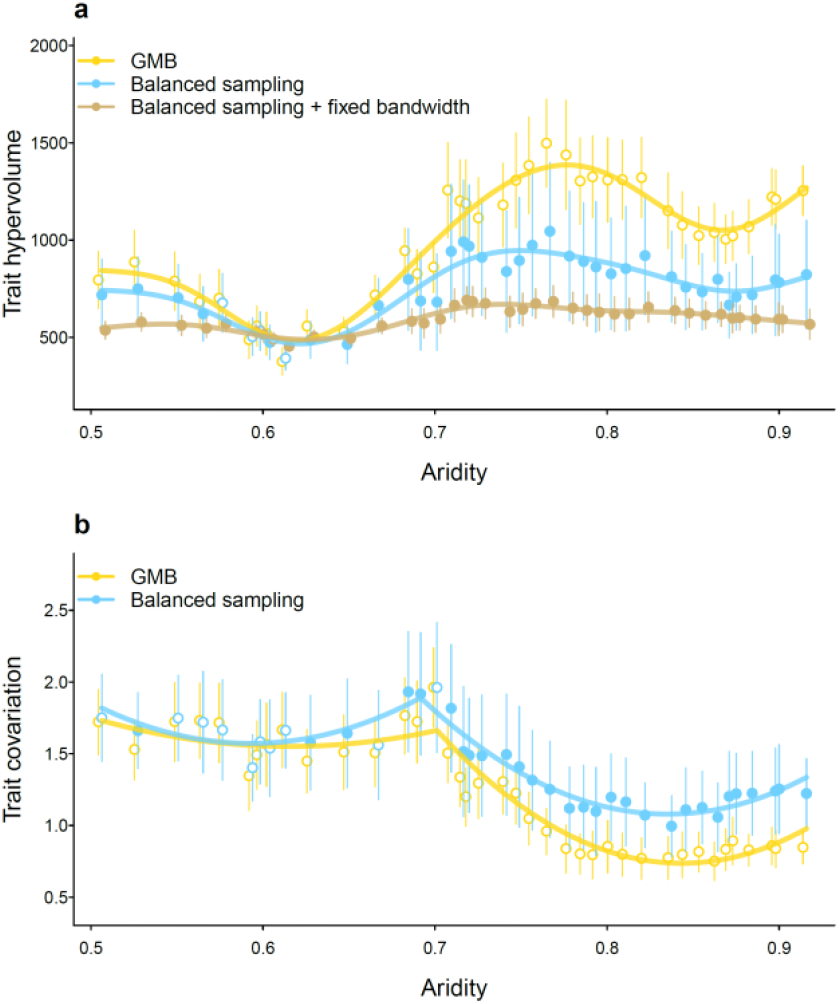
Trait hypervolume and trait integration along the aridity gradient after correcting for sampling unevenness. **a,b**, Patterns of trait hypervolume size (**a**) and trait covariation (**b**) along the aridity gradient are weaker than those presented in Gross et al.^1^ when the effects of uneven sampling and variable bandwidth are corrected. Yellow lines represent the results reported by Gross et al. (2024), while blue lines represent reanalyses correcting for unbalanced sampling by fixing the number of sites, and the brown line represents reanalyses additionally fixing the bandwidth (hypervolume only). Filled points indicate aridity intervals where the reanalysis results are significantly different from those of Gross et al. (t-test, n = 100 bootstrap samples, p < 0.01). Correcting for sampling effort and bandwidth reduced hypervolume size overall and eliminated the sharp threshold reported by Gross et al., favouring a smoother trend modelled by a GAM (Extended Data Table 1). For trait covariation, the M22 model remained the best fit (Extended Data Table 1), but the differences above and below the threshold were substantially smaller than those described in GMB.

Another critical issue in GMB is the use of adaptive bandwidths for trait hypervolume estimation^10^. The bandwidth size was determined using the Silverman estimator, which depends on the standard deviation of traits among individuals. Because GMB fixed the number of observations to 100 within each bootstrap sample, the bandwidth is exclusively driven by trait variability within each sample. However, this variability is strongly influenced by the clustering effect associated with species richness: in site-poor intervals, where observations are drawn from fewer species, individuals cluster more tightly in the trait space, leading to lower standard deviations, smaller bandwidths and, consequently, smaller hypervolumes. As a result, the correlations in GMB between hypervolume size on one hand, and bandwidth size, and the numbers of sites and individual plants in each interval on the other, are consistently high (Pearson correlations 0.95, 0.75, and 0.86, respectively, p < 0.001 in all cases; Extended Data Fig. 1). This conflation of uneven sampling intensity and bandwidth size results in a bias that overestimates the phenotypic diversity of site-rich intervals compared to site-poor ones. However, this is an issue that can be mitigated by combining the use of a single, fixed bandwidth^6,11,12^ with balanced sampling.

To better understand patterns of functional structure across the aridity gradient, we reanalysed GMB’s data, introducing a procedure to correct for sampling unevenness. Following GMB, we divided the aridity gradient into intervals and analysed trait hypervolume and integration within each interval using a bootstrapping procedure. Our reanalysis corrects the unbalanced sampling by fixing the number of sites in each bootstrap sample to eight, the minimum number found across intervals.

Additionally, to explore the effect of bandwidth, we performed hypervolume analyses using both adaptive and fixed bandwidths.

Our results revealed substantially smaller differences in phenotypic diversity at both ends of the aridity gradient than those reported by GMB. The effect of correcting the unbalanced sampling design was particularly pronounced in the most arid intervals. GAM consistently provided the best fit model of variation in hypervolume across the aridity gradient (Extended Data Table 1), challenging the existence of an aridity threshold. Standardizing the sampling design halved the difference in hypervolume between the two sides of the gradient (41.7% higher hypervolume when aridity > 0.7, vs 88.1% reported by GMB), while fixing the bandwidth further minimized this difference (17.9%; Fig. 2a). For trait integration, in agreement with GMB, the best model was the M22, which estimated a threshold at an aridity value of 0.7 (Fig. 2b). However, our reanalysis revealed a more moderate decrease in integration beyond this threshold, highlighting substantially smaller differences in phenotypic diversity than reported in GMB (24% higher integration when aridity < 0.7 vs 41% in GMB).

Our findings suggest that GMB’s claim of substantially higher phenotypic diversity in arid regions should be reconsidered in the light of methodological limitations. Our reanalysis indicates that their findings may largely reflect an uneven sampling design rather than a genuine ecological pattern.

Consequently, interpretations linking phenotypic diversity to a breakdown in environmental filtering under extreme aridity should be approached with caution. Misinterpreting these patterns could lead to oversimplified conclusions about resilience mechanisms in dryland ecosystems and the role of environmental filtering in structuring functional diversity.

We recommend that future studies carefully balance sampling effort across environmental gradients to avoid artifacts introduced by overrepresented regions. Importantly, pooling individual observations without accounting for species identity can skew estimates of trait diversity and integration due to intraspecific trait clustering. Since species identities were not available in the published dataset^13^, it was not possible for us to further explore this aspect. However, incorporating them in future analyses is essential to accurately capture intraspecific variability^9^ and prevent pseudoreplication. Future studies should explore this avenue to effectively characterize pure changes in functional structure independent from species richness patterns.

Despite these limitations, GMB represents an extraordinary collaborative effort that, by including the elementome, expands our understanding of functional diversity in ways that few studies have achieved. Importantly, while the increase in phenotypic diversity in the most arid areas may be weaker than initially presented, it remains detectable and clear—a notable result that challenges conventional expectations^14,15^. Their ambitious approach sets a valuable foundation for future research on trait diversity in response to environmental changes.

## Author contributions

All authors designed and performed the analyses. CPC and ET wrote the first draft, and all authors contributed with article writing and interpretation of results.

## Acknowledgements

We are grateful to John Davison for his comments and suggestions. This study was supported by the Estonian Ministry of Education and Research through the Estonian Research Council (PRG2142), the European Union (ERC, PLECTRUM, 101126117). GP was supported the grant IJC2020-043331-I funded by MICIU/AEI/10.13039/501100011033, “European Union NextGeneration EU/PRTR”, and by the grant PID2021-122214NA-I00 funded by MICIU/AEI/10.13039/501100011033 and by ERDF/EU.

## Data availability statement

All the data used in this paper comes from Gross et al. (2024), https://doi.org/10.57745/SFCXOO

## Code availability statement

The data and code needed to reproduce the analyses are available in Figshare (https://figshare.com/s/8e38bb66a8a9a86d652f).

## Supplementary methods

### Reconstruction of the dataset and calculation of the trait space

We performed a reanalysis of Gross et al.^1^ (hereafter ‘GMB’) quantifying trait dimensionality and hypervolume along an aridity gradient. We used the GMB dataset^2^ reporting information on 1,347 individuals and 20 trait measurements. Since the published dataset does not directly contain information on the site, we identified the site in which each observation was present by leveraging the provided metadata. To do this, we combined information on the level of aridity and the plant cover, which resulted in a univocal match between observations and sites that returned the same number of sites reported in GMB (*n* = 98, average number of sites = 19, minimum = 8, maximum = 32). For trait data manipulation and preparation, we followed exactly the script and instructions provided in GMB; in particular, we applied the same transformation and scaling procedures to all traits and used the ‘psych’ R package^3^ to calculate a principal component analysis plus varimax rotation of the first five principal components, as indicated by Horn’s parallel analysis implemented in the ‘paran’ R package^4^.

### Theoretical relationship between species richness, and trait hypervolume and covariation

Due to the clustering of conspecifics in the trait space, we expected trait hypervolume and covariation to be strongly dependent on species richness^5–7^. Because this relationship depends more strongly on species than on the number of observations, we did not expect it to be corrected by fixing the number of individual plants that are considered in trait structure estimations as done in GMB (see Main text for rationale). To demonstrate this, we created a virtual dataset of 301 species and 20 traits (the same number of species and traits as in GMB). We estimated a central coordinate in the 20-dimension trait space for each species, drawing from a multivariate Gaussian distribution based on the variance-covariance matrix of the scaled trats (i.e. all traits have unit variance and retain the covariance structure among traits). Then, for each species, we created 50 individuals around these central coordinates, following a similar procedure, but using variance = 0.1 and 0 covariance. This procedure resulted in a virtual dataset containing 15,050 individuals from 301 species, representing the idea that observations of the same species are clustered within a restricted area of the trait space^8–14^. We then performed a PCA on the scaled trait matrix and retained only the first five principal components for estimations of trait hypervolume.

Then, we simulated assemblages with a fixed number of individuals but varying levels of species richness, ranging from two to ten species. We randomly selected as many species as specified by each level of species richness, subset the virtual dataset to include only individuals from the selected number of species, and performed a bootstrap resampling with 100 individuals, similar to what was done in GMB^1^. With these 100 individuals, we estimated trait covariation (using the 20 traits) and trait hypervolume (based on the scores of individuals in the first five principal components of a PCA based on the scaled traits), following the procedure described in GMB. We repeated this procedure 100 times for each level of species richness. As expected, these simulations revealed a strong dependency between species richness, hypervolume size and trait integration (see Fig. 1 in the main text).

### Trait integration and hypervolume calculation

To evaluate how trait integration and hypervolume size varied along the aridity gradient, we implemented the same sliding-window procedure presented in GMB, with some slight adjustments to correct for unbalanced sampling. We divided the aridity gradient into the same 39 intervals used in GMB and compared our integration and hypervolume results with the ones published in GMB^2^.

While the relationship between hypervolume size and trait covariation depends ultimately on species richness, species identities were not available in the published dataset^2^, so it was not possible to further explore this aspect. However, as explained in the main text, the imbalance in site number across intervals likely results in a similar unbalance in species richness. To further show this, we explored GMB’s data and calculated Pearson’s correlations among hypervolume size, the number of sites and the number of individual plants included in each aridity interval. These correlations revealed an intrinsic dependency between the dimensionality of the trait space, the size of the hypervolume and the number of sites considered along the aridity gradient (Extended Data Fig. 1, see below for methodological details).

To address the unbalanced sampling design in GMB, we examined the variation in the number of sites across aridity intervals. The interval with the fewest sites contained eight, so we opted to simulate a sampling scheme where eight sites were sampled per aridity interval to achieve a more balanced approach for estimating functional trait structure. Following this strategy, we replicated the bootstrap analyses performed in GMB, but within each aridity interval, we restricted the number of sites that were considered to eight (hereafter referred to as ‘balanced sampling’). For each bootstrap sample, we first randomly selected eight sites without replacement from all sites within the aridity interval. From these selected sites, we then bootstrapped 100 observations, ensuring that each bootstrap sample consistently included data from all eight sites (i.e., discarding bootstrap samples where observations originated from fewer than eight sites). This process was repeated 100 times for each aridity interval. Using these samples, we estimated trait hypervolume and integration following the same procedure described in GMB. Specifically, trait integration was estimated as the variance of the adjusted eigenvalues after parallel analysis^4^ based on the 20 traits, while trait hypervolume was calculated using multivariate Gaussian kernel density estimation as implemented in the R package ‘hypervolume’^15^ considering the five-dimensional varimax-rotated space described in GMB.

The bandwidth in the Gaussian kernel density estimation procedure controls the amount of ‘padding’ around each observation^13,14^, effectively controlling the influence of each data point on the density estimate. Since bandwidth fundamentally affects the structure of the resulting hypervolume, comparing hypervolumes estimated with different bandwidths can lead to inconsistencies: larger bandwidths lead to larger hypervolume sizes. To ensure fair and meaningful comparisons, a frequent practice in studies analysing the sizes of trait hypervolumes is to standardize the analyses by using a fixed bandwidth ^5,16–19^. However, GMB applied an adaptive bandwidth approach, where bandwidths were adjusted for each bootstrap sample and across the aridity gradient, potentially introducing variability that complicates direct comparisons.

To examine the potential spurious relationship between bandwidth and hypervolume size in GMB, we replicated their hypervolume estimation procedure following the provided scripts^2^. We stored the bandwidth vector for the hypervolume estimation of each bootstrap sample—formed by the individual bandwidths of the five dimensions of the varimax-rotated PCA space— and calculated the product of its components. The resulting product (from now on referred to as ‘bandwidth size’) reflects the amount of padding applied around each individual observation. Then, we examined how bandwidth size changed along the aridity gradient, as well as its relationship to the hypervolume size values reported in GMB (i.e., with unbalanced sampling), and the number of sites and total observations present in each aridity interval. Our results indicate an extremely high correlation between bandwidth size and hypervolume (Extended Data Fig. 1), calling into question the validity of the adaptive bandwidth approach to compare samples along the aridity gradient.

Finally, we performed two hypervolume estimations for each bootstrap sample in the balanced sampling approach, using two alternative bandwidth strategies. First, we replicated the adaptive bandwidth estimation used in GMB, where a bandwidth vector was estimated for each bootstrap sample using the Silverman estimator from the hypervolume package^15^ (‘Balanced sampling’ in Fig. 2a) Second, we estimated hypervolumes using a fixed bandwidth (‘Balanced sampling + fixed bandwidth’ in Fig. 2a). To select the fixed bandwidth, we first applied the Silverman estimator to all individual observations within each aridity interval (39 bandwidth vectors in total) and then averaged these vectors across intervals. This procedure provides a compromise bandwidth representative of the entire dataset, ensuring consistent smoothing across bootstrap samples.

To assess the robustness of hypervolume estimates under different fixed bandwidths, we performed a sensitivity analysis. We adjusted the fixed bandwidth for each principal component by ± 50% to create three different fixed bandwidth scenarios: fixed (the average bandwidth described above), fixed_High (+50%) and fixed_Low (-50%). Hypervolumes were then calculated for each scenario using 25 bootstrap repetitions per aridity interval. The sensitivity analysis showed that the choice of one or another fixed bandwidth does not affect the hypervolume pattern along the aridity gradient, with there were high correlations between the hypervolumes estimated using the fixed bandwidth and those obtained using the fixed_High and fixed_Low alternatives (Pearson correlation: fixed vs. fixed_High = 0.95; fixed vs. fixed_Low = 0.85; p < 0.001 for both). Because of this robustness, we report only the values obtained using the fixed bandwidth approach.

### Data analysis

We compared the results of trait hypervolume and integration provided by GMB with the different corrections explained in the previous section (balanced sampling for integration, and balanced sampling and balanced sampling + fixed bandwidth for hypervolume). We assessed the responses of trait integration and hypervolume along the aridity gradient using the same set of models used in GMB. Specifically, we fitted seven models: a linear model, five threshold models (step, segmented, stegmented, M12, M22) implemented using the R package ‘chngpt’^20^, and a generalized additive model (GAM) using the ‘mgcv’ package^21^. Model selection was done using the Akaike (AIC) and Bayesian Information Criteria (BIC). Finally, within each aridity interval, we performed a t-test to identify the levels at which the reanalysis results are significantly different from those of GMB (n = 100 bootstrap samples).

## Extended Data

**Extended Data Table 1.**
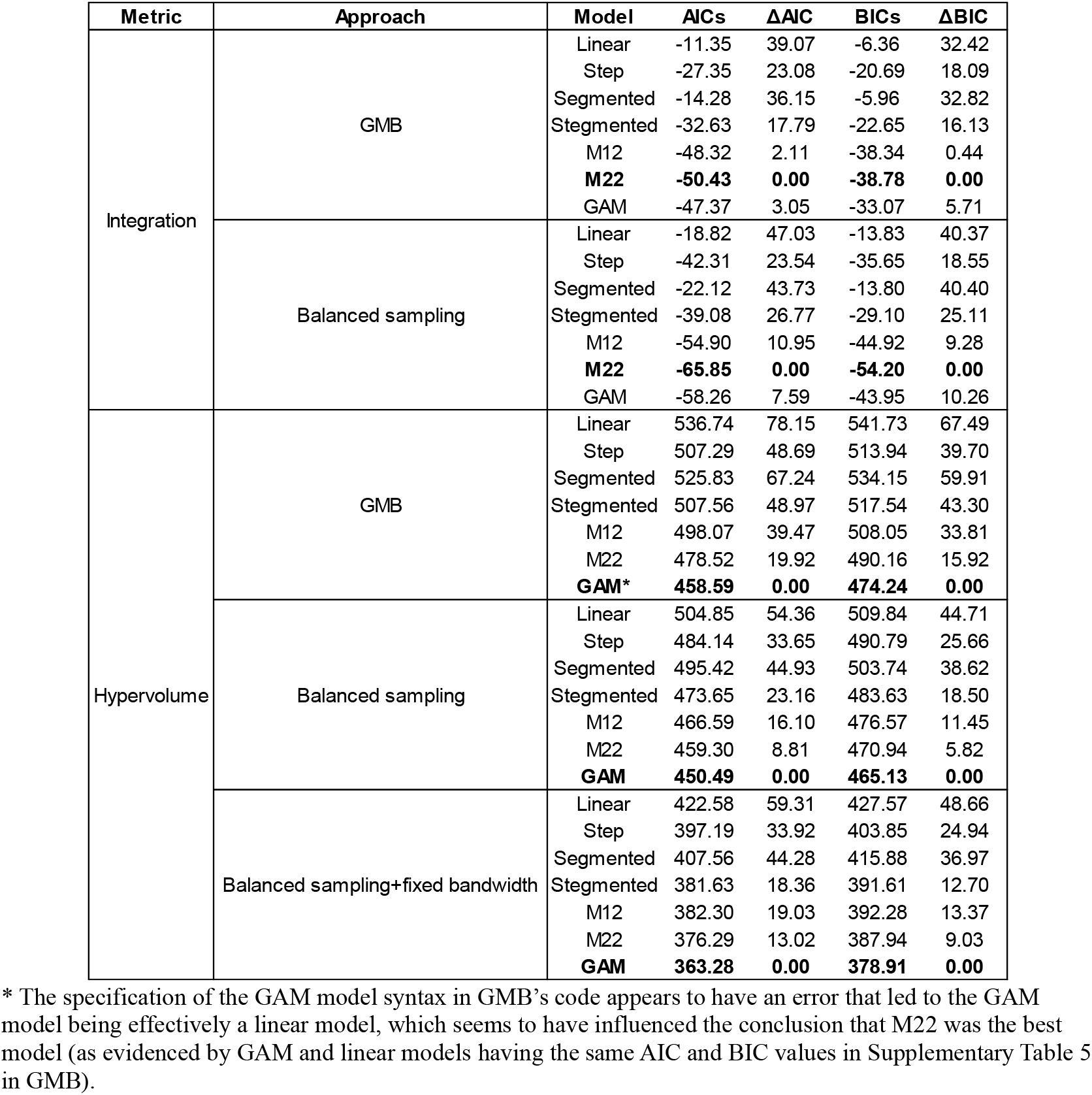
Performance of the models used to evaluate the response of trait integration and hypervolume to aridity. For each model, we provided the Akaike Information Criterion (AIC) and Bayesian Information Criterion (BIC) along with the difference with the best model. The best model is highlighted in bold. “GMB” indicate the approach used in Gross et al., “Balanced sampling” refers to the reanalyses correcting for unbalanced sampling by fixing the number of sites within aridity intervals, and the “Balanced sampling+fixed bandwidth” represents reanalyses additionally fixing the bandwidth (hypervolume only).

| Metric | Approach | Model | AICs | ΔAIC | BICs | ΔBIC |
| --- | --- | --- | --- | --- | --- | --- |
| Integration | GMB | Linear | -11.35 | 39.07 | -6.36 | 32.42 |
|  |  | Step | -27.35 | 23.08 | -20.69 | 18.09 |
|  |  | Segmented | -14.28 | 36.15 | -5.96 | 32.82 |
|  |  | Stegmented | -32.63 | 17.79 | -22.65 | 16.13 |
|  |  | M12 | -48.32 | 2.11 | -38.34 | 0.44 |
|  |  | **M22** | **-50.43** | **0.00** | **-38.78** | **0.00** |
|  |  | GAM | -47.37 | 3.05 | -33.07 | 5.71 |
|  | Balanced sampling | Linear | -18.82 | 47.03 | -13.83 | 40.37 |
|  |  | Step | -42.31 | 23.54 | -35.65 | 18.55 |
|  |  | Segmented | -22.12 | 43.73 | -13.80 | 40.40 |
|  |  | Stegmented | -39.08 | 26.77 | -29.10 | 25.11 |
|  |  | M12 | -54.90 | 10.95 | -44.92 | 9.28 |
|  |  | **M22** | **-65.85** | **0.00** | **-54.20** | **0.00** |
|  |  | GAM | -58.26 | 7.59 | -43.95 | 10.26 |
| Hypervolume | GMB | Linear | 536.74 | 78.15 | 541.73 | 67.49 |
|  |  | Step | 507.29 | 48.69 | 513.94 | 39.70 |
|  |  | Segmented | 525.83 | 67.24 | 534.15 | 59.91 |
|  |  | Stegmented | 507.56 | 48.97 | 517.54 | 43.30 |
|  |  | M12 | 498.07 | 39.47 | 508.05 | 33.81 |
|  |  | M22 | 478.52 | 19.92 | 490.16 | 15.92 |
|  |  | **GAM\*** | **458.59** | **0.00** | **474.24** | **0.00** |
|  | Balanced sampling | Linear | 504.85 | 54.36 | 509.84 | 44.71 |
|  |  | Step | 484.14 | 33.65 | 490.79 | 25.66 |
|  |  | Segmented | 495.42 | 44.93 | 503.74 | 38.62 |
|  |  | Stegmented | 473.65 | 23.16 | 483.63 | 18.50 |
|  |  | M12 | 466.59 | 16.10 | 476.57 | 11.45 |
|  |  | M22 | 459.30 | 8.81 | 470.94 | 5.82 |
|  |  | **GAM** | **450.49** | **0.00** | **465.13** | **0.00** |
|  | Balanced sampling+fixed bandwidth | Linear | 422.58 | 59.31 | 427.57 | 48.66 |
|  |  | Step | 397.19 | 33.92 | 403.85 | 24.94 |
|  |  | Segmented | 407.56 | 44.28 | 415.88 | 36.97 |
|  |  | Stegmented | 381.63 | 18.36 | 391.61 | 12.70 |
|  |  | M12 | 382.30 | 19.03 | 392.28 | 13.37 |
|  |  | M22 | 376.29 | 13.02 | 387.94 | 9.03 |
|  |  | **GAM** | **363.28** | **0.00** | **378.91** | **0.00** |

\* The specification of the GAM model syntax in GMB’s code appears to have an error that led to the GAM model being effectively a linear model, which seems to have influenced the conclusion that M22 was the best model (as evidenced by GAM and linear models having the same AIC and BIC values in Supplementary Table 5 in GMB).

**Extended Data Figure 1.**
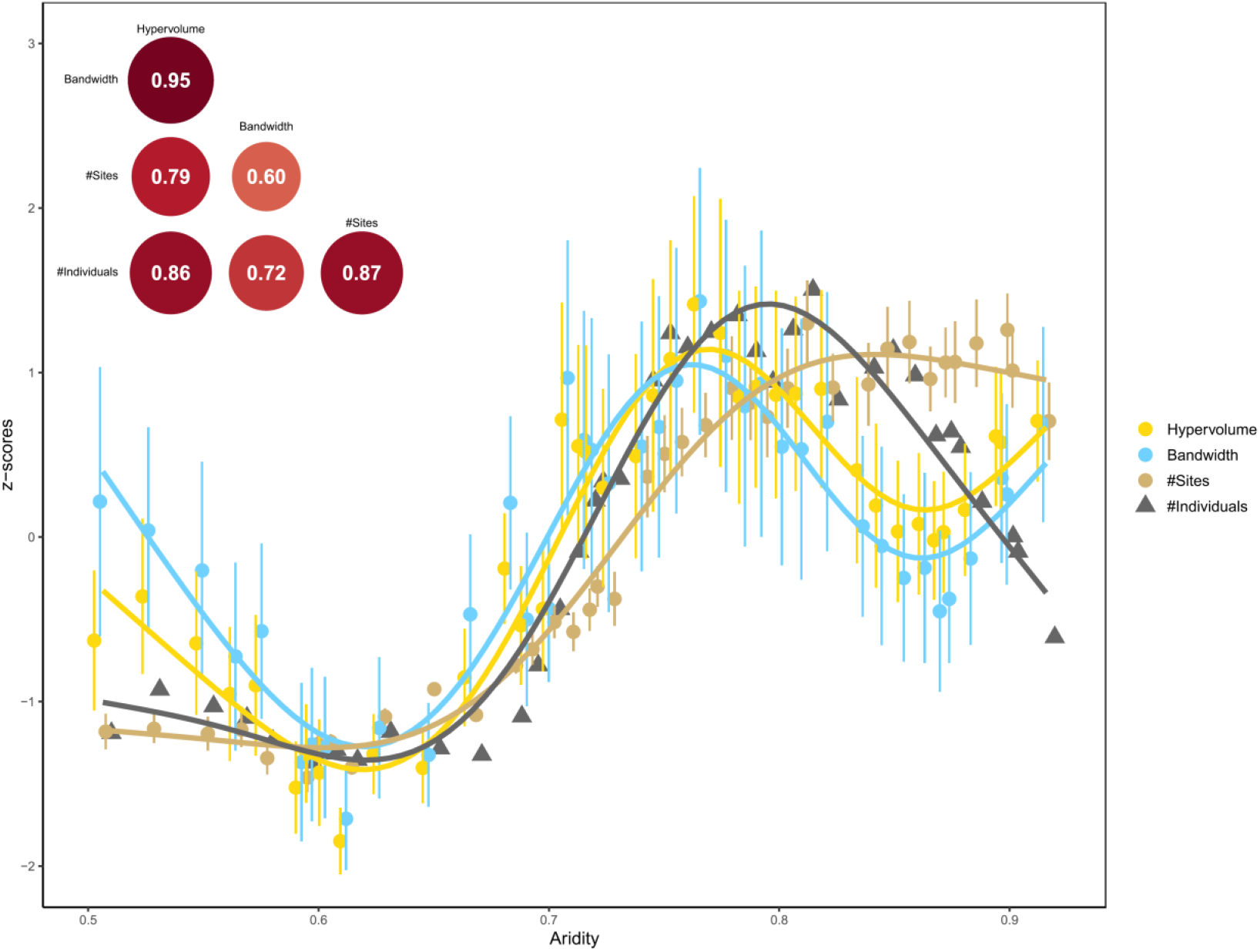
Variation of trait hypervolume size as calculated in GMB (yellow), size of the bandwidth (blue), number of different sites included in each bootstrap sample (brown) and the total number of individuals in each aridity interval (grey) along the aridity gradient. All values were scaled to facilitate visual comparisons. Points represent the mean value within each aridity interval while the interval lines indicate one SD (except for number of individuals, which do not vary within aridity intervals). Trend lines represent a GAM model computed on the mean values per window as a function of the aridity level. The upper inset indicates the pairwise Pearson’s correlations of the mean values in the aridity windows.

**Extended Data Figure 2.**
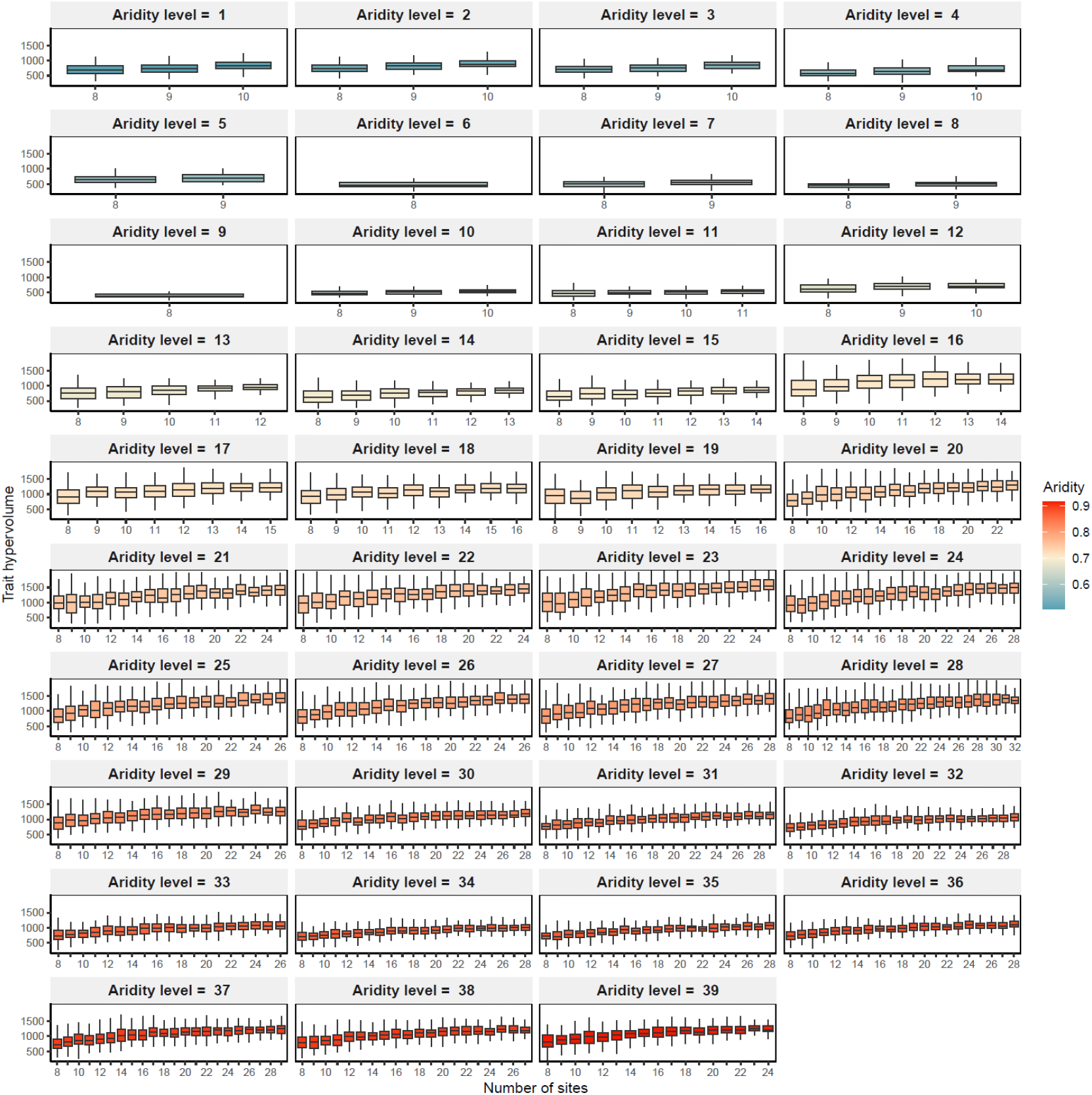
Variation of trait hypervolumes calculated with adaptive bandwidth as a function of the number of sites within each aridity interval. For each interval, bootstrap samples were restricted to include different numbers of sites (ranging from eight to the total number of sites available in the interval), following the balanced sampling procedure. Each bootstrap sample included 100 observations. Results demonstrate an increasing relationship between the number of sites considered in each aridity level and hypervolume size, highlighting that values of hypervolumes across aridity levels with varying site numbers are not directly comparable.

**Extended Data Figure 3.**
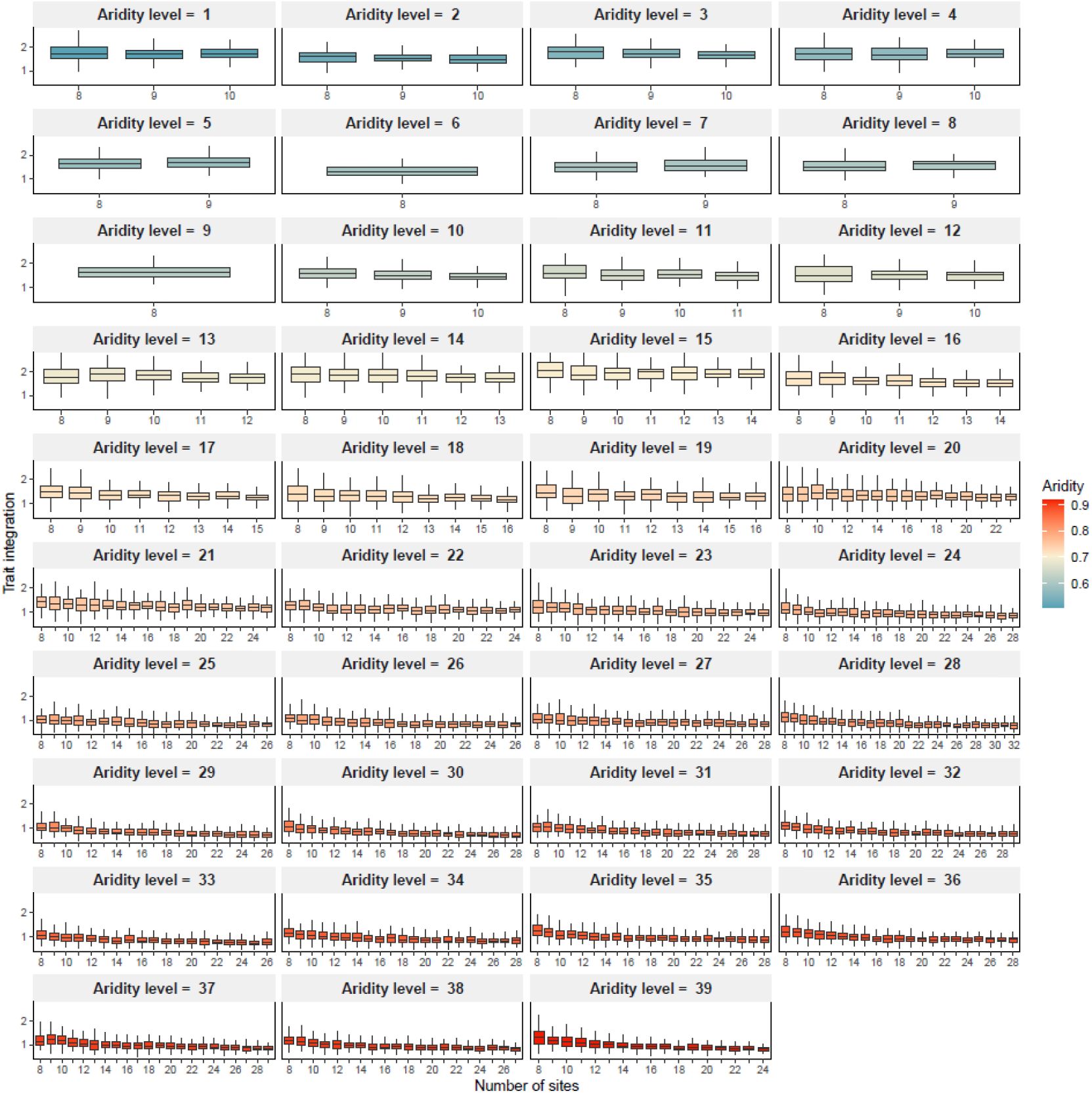
Variation of trait integration calculated with adaptive bandwidth as a function of the number of sites within each aridity interval. For each interval, bootstrap samples were restricted to include different numbers of sites (ranging from eight to the total number of sites available in the interval), following the balanced sampling procedure. Each bootstrap sample included 100 observations. Results demonstrate a decreasing relationship between the number of sites considered in each aridity level and trait integration, highlighting that values of trait integration across aridity levels with varying site numbers are not directly comparable.

